# Impact of Human Behavioral Papers at Journal of Neuroscience

**DOI:** 10.1101/248526

**Authors:** David Herzfeld, Reza Shadmehr

**Affiliations:** Laboratory for Computational Motor Control, Department of Biomedical Engineering, Johns Hopkins School of Medicine

## Abstract

A recent policy change at Journal of Neuroscience (JN) has significantly increased editorial “desk rejections”, reducing the number of papers that are considered for publication. Survey results from 130 scientists suggested that the new policy may have had a particularly large impact on studies that focused on human behavioral techniques. To quantify the effects of the new policy, we gathered data on all papers ever published in JN (~35,000), as well as all papers that had cited the JN papers (~2.7 million papers). We found that the recent change in editorial policy had disproportionately affected rejection rates of human behavioral papers: since 2015, the number of human behavioral papers as a proportion of all papers published in JN has seen a 30% decline. While there has been a long-term declining trend in the journal’s impact factor, we found that this declining impact factor was shared by both human behavioral papers as well as other papers in the journal. This suggested that whatever may have been the source of the declining impact factor at JN, this source was affecting the various areas of research equally. That is, it was unlikely that papers in any one field were responsible for the declining impact factor number at JN. Indeed, when impact was measured over the long-term, we found that the average human behavioral paper at JN consistently outperformed other papers, generating a significantly higher number of citations per year at 5 and 10 years post publication.

## Introduction

There is a trend among some neuroscience journals to increase their focus on research that rely on invasive techniques to describe neural function, and reduce the number of papers that use non-invasive, behavioral techniques that only by inference, describe neural function (1). For example, in the January 6, 2016 issue of *Journal of Neuroscience* (JN), the editors outlined a new policy that aimed to increase the rate of editorial rejections of submitted manuscripts (2). In a follow-up article that was published in the June 1, 2016 issue, the editors further clarified the new policies (3). They provided examples of the kind of research that they would not consider for review: “…purely biophysical or behavioral studies should provide novel insights into, and make specific predictions about, neural mechanisms or neural representations.” Because the policy statement focused on behavioral experiments in particular, we conducted a survey to gauge the impact of this policy on neuroscientists who submit papers to JN. We then followed up the survey with quantitative analysis of the impact of human behavioral papers, published over the entire history of the journal.

We performed this analysis in order to answer a simple question: what has been the long-term impact of papers that have focused on non-invasive, behavioral techniques, as compared to other papers at the JN? Here, we report on the survey results, and the results of the impact factor analysis.

## Results

In October of 2017, 130 scientists, representing authors of 342 papers in JN, responded with examples of editorial rejection letters that they had received from JN. They reported 148 desk-rejections. In approximately 75% of the cases, the editors felt that the work “lacked insights into neural mechanisms.” The letters described the reasons for editorial rejection:

- “For purely behavioral studies, our criteria have evolved to require that a behavioral study provides novel insights into the underlying neural representations and mechanisms.”
- “The study does make some experimental predictions, but they are all behavioral.”
- “while the study had a number of strengths, the emphasis was on behavioral processes rather than neural mechanisms which are the focus for The Journal of Neuroscience.”
- “It is, unfortunately, the case that it is not possible for us to consider manuscripts where the emphasis is on behavior.”
- “I am afraid that it has become much rarer for behavior only manuscripts to be sent out for review at The Journal of Neuroscience.”

We followed this survey with analysis of historical data. We collected information on all papers published in the history of JN (approximately 35,000 papers), as well as information on all papers that had cited these papers (approximately 2.7 million papers, citation data was collected from Web of Science, see Methods). Our aim was to quantify how the new policy had affected the ability of scientists that relied primarily on behavioral measures to publish their results in JN. However, because the change in editorial policy appeared to have come about because of a two-decade long decline in the impact factor of the journal, we also quantified the citation impact of human behavioral papers at JN as compared to all other papers at the journal.

From 1981 (inception of the journal) to 2011, the number of papers published per year increased at a fairly constant rate (Fig. 1A, an increase of approximately 54 papers/year). However, since 2013 the number of papers published per year has seen a decline. We searched the abstract and title of each JN paper for keywords that included “human”, and excluded words such as “EEG”, “fMRI”, etc. That is, we looked for papers that could be classified as behavioral, with little or no inclusion of neural correlates of behavior (see Table 1). We labeled these as “human behavioral” papers, resulting in about 1,200 papers, representing 4% of papers published at JN.

**Figure 1.**
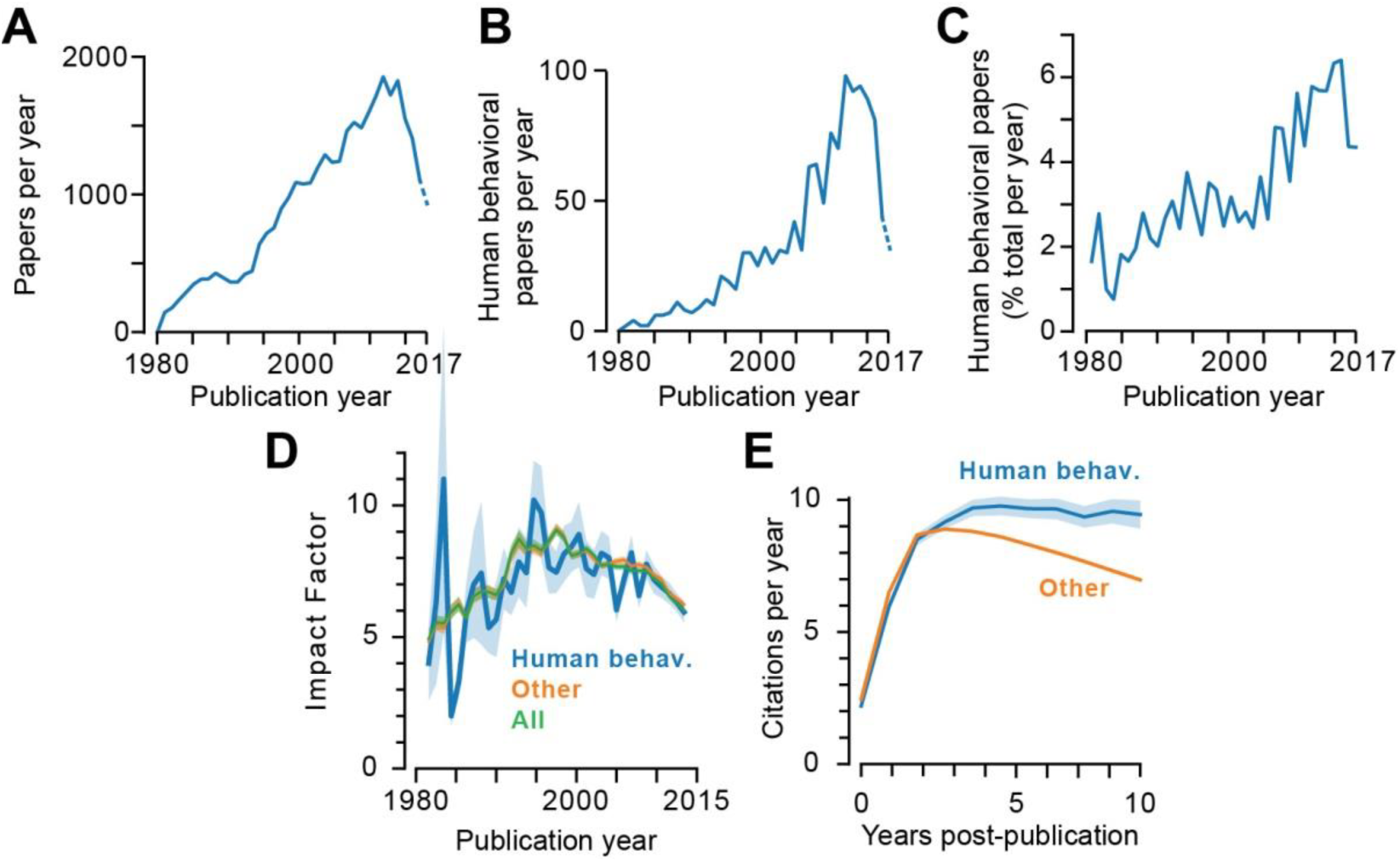
Analysis of papers published in *Journal of Neuroscience*. **A**. Number of papers published per year. **B**. Human behavioral papers published per year. **C**. Human behavioral papers as a percent of total papers published each year. **D**. Impact Factor (IF) of human behavioral papers, other papers, and all papers in JN. IF counts the number of citations generated by the paper in the year of publication and the year following publication. Shaded regions are SEM. **E**. Citation rate of human behavioral papers and other papers at JN. Shaded regions are SEM.

**Table 1.**
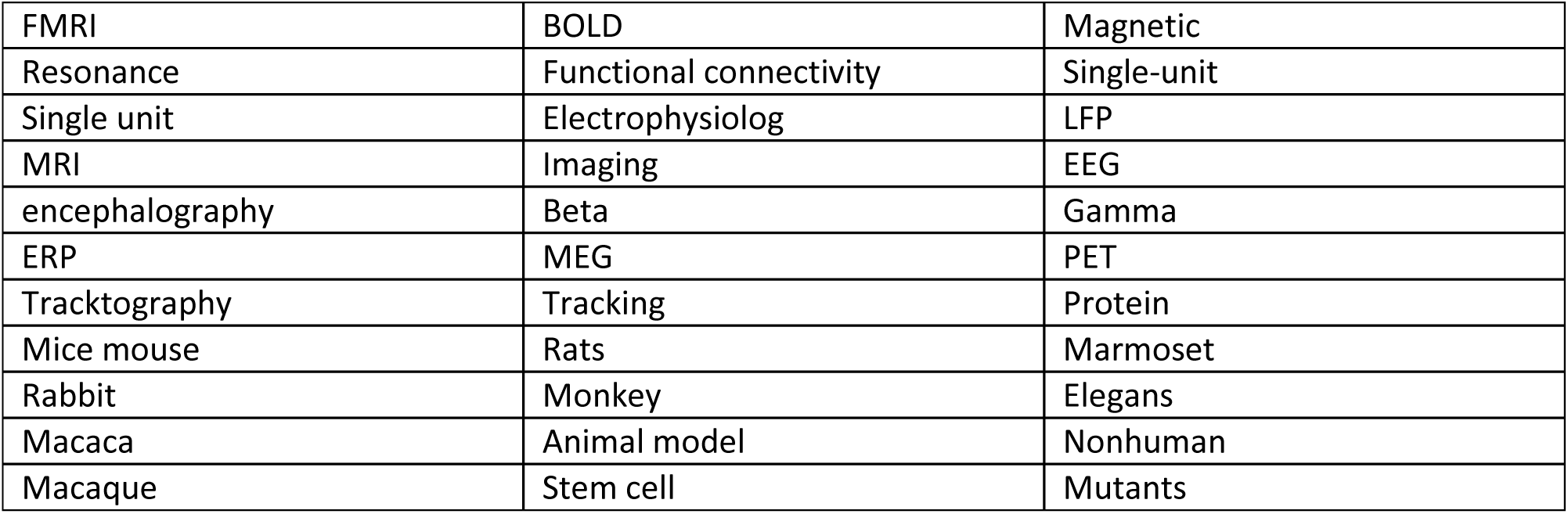
List of keywords used to exclude non-human behavioral studies. Journal publications were considered as a non-human study if the title or abstract contained one of more of these (caseinsensitive) keywords. Each word began and ended with ‘*’, allowing partial matches to words in the abstract or title. These ‘*’ are omitted for clarity.

|  |  |  |
| --- | --- | --- |
| FMRI | BOLD | Magnetic |
| Resonance | Functional connectivity | Single-unit |
| Single unit | Electrophysiolog | LFP |
| MRI | Imaging | EEG |
| encephalography | Beta | Gamma |
| ERP | MEG | PET |
| Tracktography | Tracking | Protein |
| Mice mouse | Rats | Marmoset |
| Rabbit | Monkey | Elegans |
| Macaca | Animal model | Nonhuman |
| Macaque | Stem cell | Mutants |

To quantify how the new policy may have affected the ability to publish human behavioral papers, we measured the human behavioral papers as a percentage of total papers published each year. Our results illustrated that while human behavioral papers have historically represented only a small minority of papers published at the journal (Fig. 1C), the percentage had gradually increased, reaching a high of around 6% in 2015. However, in 2016 (the inception of the new policy) this percentage saw its largest decline in the journal’s history (a reduction of roughly 30%). Data for 2017 (collected until November 2017) confirmed that the reduced rates have persisted. Therefore, the data suggest that the policy that went into effect in 2016 has had a disproportionately negative effect on number of human behavioral studies published in JN.

We next quantified the impact factor of every paper published in JN. The Journal Impact Factor (JIF) is a metric that describes the average impact factor across all papers in the journal over the 0-1 year calendar period following publication. JIF of JN exhibited a gradual increase from inception of the journal, reaching a peak in 1998 (Fig. 1D). However, since 1998 the JIF has shown a gradual decline. We computed the impact factor for human behavioral papers separately and found that as a group, they tracked the JIF. This co-variance between impact factor of human behavioral papers and all other papers is noteworthy because of the fact that human behavioral papers represent at most 6% of the papers published annually at the journal. The presence of a long-term co-variance suggests that there exists a common factor that is influencing the quality of the published papers, regardless of behavioral or otherwise.

How does the impact of human behavioral papers compare to other papers published at the journal? We measured the number of citations generated by a human behavioral paper as a function of years since its publication, and compared it to citation history of other papers at JN (Fig. 1E). We found that human behavioral papers, like other papers, increased their citation rate in the 0-3 year period following publication. However, whereas for human behavioral papers the citation rate reached about 10/year in the 4^th^ year and was sustained for up to 10 years, for other papers in JN the citation rate peaked in the 3^rd^ year and then declined. As a result, a typical human behavioral paper produced significantly more citations over a 10 year period than other papers in the journal (Wilcoxon rank sum, p=0.009). This suggests that the long-term impact of a human behavioral paper at JN is significantly greater than other papers.

## Discussion

We found evidence suggesting that the 2016 change in editorial policies at JN appears to be disproportionately affecting rejection rates of human behavioral papers: as measured against the year preceding the installment of the policy, the number of human behavioral papers as a proportion of all papers published in JN has seen a 30% decline.

While there has been a long-term trend in the journal’s impact factor, we found that this declining impact factor is shared by both human behavioral papers as well as other papers in the journal. This suggests that whatever may be the source of the declining impact factor of the journal; this source is affecting the various types of papers equally, making it unlikely that any one field is responsible for the declining numbers. Indeed, when impact was measured over the long-term, we found that the average human behavioral paper consistently outperformed other papers, generating a significantly higher number of citations per year at 5 and 10 years post publication.

In a thoughtful perspective, (1) considered the question of whether behavioral experiments are fundamental to advancing neuroscience. They noted that whereas the focus of neuroscience appears to have shifted to neural circuits, behavioral experiments provide the guiding vision of what that circuit might be doing. They wrote: "when scientists ask 'how does the brain generate behavior,' they are in fact asking a question best approached through behavioral work, specifically task analysis, aided by theory, that allows behavior to be decomposed into separable modules and processing operations … The neural basis of behavior cannot be properly characterized without first allowing for independent, detailed study of the behavior itself."

History of science provides vivid examples where entire fields start with experiments that probe nature with quantitative, but non-invasive measures. Gregor Mendel’s experiments on traits of peas were published in 1866, and cited only 3 times in the next 35 years. In it, he meticulously measured the statistical nature of heredity, and of course, made no mention of the molecular basis of the process. Today, experiments that use non-invasive behavioral measures may make fundamental observations about the function of the brain, but there is a danger that because of the current policies, results of these experiments may not be published in the *Journal of Neuroscience* because of the editorial insistence that experiments must provide insights into neural mechanisms.

## Methods

We analyzed the citation history of 34,756 papers published in the *Journal of Neuroscience* (JN) from inception of the journal in 1981 to Oct. 20, 2017. We specifically analyzed only the data for publications index in PubMed as “Journal Articles,” intentionally removing review articles and commentaries from analysis. We downloaded the meta-data for each publication from the PubMed database, including title, full-text abstract, publication date, volume, issue, DOI, etc. We excluded papers from analysis where this meta-data was incomplete. Typically, these were manuscripts without full text abstracts, published in the early years following JN inception. Our final dataset consisted of 31,693 research articles with complete meta-data.

Our goal was to determine the impact factor (IF) and average citations per year for each paper published in JN. In order to compute the IF, we required the complete citation record for each of the 31,693 papers we identified from the PubMed dataset. The official impact factor for each journal is reported by Thomas Reuters using their Web of Science citation database. To allow direct comparison with the IF as reported by Reuters, we similarly used their Web of Science database to determine the total number of citations for each paper as a function of time. We obtained Johns Hopkins institutional access to the Web of Science database and, for each JN manuscript, downloaded meta-data for all publications indexed in Web of Science that cited each manuscript. Specifically, we downloaded approximately the date of publication for 2.7 million publications that cited the research articles published in JN. These citations must have been indexed by Web of Science prior to Oct. 20, 2017.

We asked whether human behavioral studies had a differential effect on the IF. To segregate human behavioral studies from all other study types, we used keyword matching in the title and abstract. To be considered a human behavioral study, the abstract or title must have contained one or more of the following case-insensitive words: “human”, “participant”, “patient”, “pyschophy*”, “subjects” (where the ‘*’ denotes a wildcard). In addition to these required keywords, the abstract and title were not allowed to contain any of the keywords shown in Table 1. These set of strict criteria led to the identification of 1,200 human behavior manuscripts.

## Reference

1. Krakauer JW, Ghazanfar AA, Gomez-Marin A, MacIver MA, and Poeppel D (2017) Neuroscience needs Behavior: correcting a reductionist bias. Neuron 93:480–490.

2. Picciotto, M. A Message from the Editor-in-Chief. J. Neurosci. 36, ii–ii (2016).

3. Journal of Neuroscience. Changes to the Review Process. J. Neurosci. 36, 5907–5908 (2016).

